# Spectral tuning of adaptation supports coding of sensory context in auditory cortex

**DOI:** 10.1101/534537

**Authors:** Mateo Lopez Espejo, Zachary P. Schwartz, Stephen V. David

## Abstract

Perception of vocalizations and other behaviorally relevant sounds requires integrating acoustic information over hundreds of milliseconds, but sound-evoked activity in auditory cortex typically has much shorter latency. It has been observed that the acoustic context, i.e., sound history, can modulate sound evoked activity. Contextual effects are attributed to modulatory phenomena, such as stimulus-specific adaption and contrast gain control. However, an encoding model that links context to natural sound processing has yet to be established. We tested whether a model in which spectrally tuned inputs undergo adaptation mimicking short-term synaptic plasticity (STP) can account for contextual effects during natural sound processing. Single-unit activity was recorded from primary auditory cortex of awake ferrets during presentation of noise with natural temporal dynamics and fully natural sounds. Encoding properties were characterized by a standard linear-nonlinear spectro-temporal receptive field model (LN STRF) and STRF variants that incorporated STP-like adaptation. In the adapting models, STP was applied either globally across all input spectral channels or locally to subsets of channels. For most neurons, STRFs incorporating local STP predicted neural activity as well or better than the LN and global STP STRF. The strength of nonlinear adaptation varied across neurons. Within neurons, adaptation was generally stronger for spectral channels with excitatory than inhibitory gain. Neurons showing improved STP model performance also tended to undergo stimulus-specific adaptation, suggesting a common mechanism for these phenomena. When STP STRFs were compared between passive and active behavior conditions, response gain often changed, but average STP parameters were stable. Thus, spectrally and temporally heterogeneous adaptation, subserved by a mechanism with STP-like dynamics, may support representation of the complex spectro-temporal patterns that comprise natural sounds across wide-ranging sensory contexts.

## Introduction

Vocalizations and other natural sounds are characterized by complex spectro-temporal patterns. Discriminating sounds like speech syllables requires integrating information about changes in frequency content over many tens to hundreds of milliseconds [1–3]. Models of sensory encoding for auditory neurons, such as the widely used linear-nonlinear spectro-temporal receptive field (LN STRF), seek to characterize sound coding generally. That is, they are designed to predict time-varying responses to any arbitrary stimulus, including natural sounds [4]. When used to study auditory cortex, these LN models typically are able to measure tuning properties only with relatively short latencies (20-50 ms), which prevents them from encoding information about stimuli over a longer period of time [5,6]. It remains an open question how the auditory system integrates spectro-temporal information from natural stimuli over longer periods.

Classic STRF models cannot account for integration over longer timescales, but studies of spectro-temporal context have shown that auditory-evoked activity can be modulated by stimuli occurring hundreds to thousands of milliseconds beforehand [7–9]. These results have generally been interpreted in the context of pop-out effects for oddball stimuli [10–13] or gain control to normalize neural activity in the steady state [14–16]. Encoding models that incorporate recurrent gain control or nonlinear adaptation have been shown to provide better characterization of auditory-evoked activity in the steady state, indicating that these properties of neurons may contribute to context-dependent coding on these longer timescales [12,17–21]. Some models have been shown to account for cortical responses to natural stimuli more accurately than the LN STRF [19,21–23], and others have been proposed that have yet to be tested with natural stimuli [18,20,24,25]. These findings suggest that an adaptation mechanism plays a central role in context-dependent coding, but there is no clear consensus on the essential components of a model that might replace the LN STRF as a standard across the field.

Short-term synaptic plasticity (STP) is a widely-observed phenomenon in nervous system. Upon sustained stimulation, the efficacy of synapses is depressed or facilitated until stimulation ceases and synaptic resources are allowed to return to baseline [26,27]. As sensory inputs are integrated by the neural processing hierarchy, synapses can undergo different degrees of plasticity, which depend on the specific pattern of stimulation. We hypothesized that nonlinear adaptation with STP-like properties may play a general role in auditory processing. While the precise mechanism producing nonlinear adaptation can take other forms than STP (e.g., feedforward inhibition, postsynaptic inhibition [11,28]), all these mechanisms support a fundamentally similar algorithm for encoding spectro-temporal features. The focus of this study is whether functional properties of auditory neurons are impacted significantly by such a mechanism at the algorithmic level and, in particular, if this adaptation occurs independently across inputs with different sound frequency. Regardless of precise mechanism, a population of neurons with spectrally tuned adaptation may support a rich code for information over the many hundreds of milliseconds required to discriminate spectro-temporally complex natural sounds [29,30].

To test for spectrally-tuned adaptation, we developed a stimulus in which two independently modulated narrowband noise stimuli were presented in the receptive field of neurons recorded in primary auditory cortex (A1) of awake ferrets. We then used the approach of comparing the prediction accuracy of neural encoding models to test for STP-like effects [4,31]. We fit an encoding model based on the STRF in which inputs adapted either locally to one spectral band or globally across all channels. For many neurons, locally tuned adaptation provided a more accurate prediction of neural activity, supporting the idea of channel-specific adaptation. The strength and tuning of adaptation was heterogeneous across the A1 population, consistent with the idea that a diversity of spectrally tuned adaptation supports a rich basis for encoding complex natural sounds. We observed the same pattern of results for models fit to a library of fully natural sounds.

We also asked how changes in behavioral state, which can influence response gain and selectivity, affected nonlinear adaptation properties in A1 [32–38]. We compared model STP parameters between passive listening and during a behavior that required detecting a tone in a natural noise stream. While the gain of the neural response could fluctuate substantially with behavioral state, STP was largely stable across behavior conditions. This finding suggests that, unlike response gain, nonlinear adaptation properties are not influenced by behavioral state and may instead be critical for stable encoding of spectro-temporal sound features [29,39]. Together, these findings demonstrate that during natural hearing, a simple, STP-like mechanism can explain many aspects of context-dependent sound coding. Moreover, these processes, typically associated with steady-state adaptation to different contexts, such as SSA, can play a more dynamic role, continuously shaping the representation of spectro-temporally complex natural sounds.

## Results

### Encoding models reveal spectrally tuned adaptation in primary auditory cortex

This study characterized how primary auditory cortex (A1) integrated information from dynamic, naturalistic stimuli over time and frequency. Data were recorded from 200 single units in A1 of 5 passively listening ferrets during presentation of two band vocalization-modulated noise (Fig. 1A-B, [17,40]). The stimulus contained complex natural temporal statistics but simple spectral properties. Thus it allowed an experimental focus on nonlinear temporal processing in the presence of multiple spectral features. Noise bands were one-quarter octave and modulated by different natural vocalization envelopes. Both bands were positioned so that they fell in the spectral receptive field of recorded neurons, as measured by briefly presented tones or noise bursts (Fig 1B).

**Figure 1.**
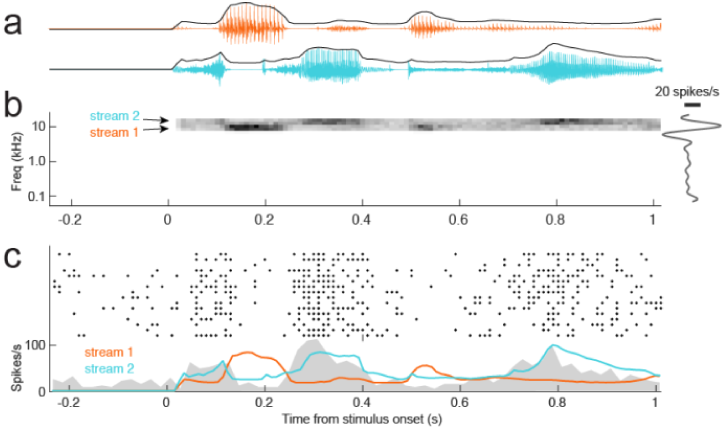
**A.** Two example natural vocalization waveforms show characteristic interspersed epochs of high sound energy and silence. Each sound has a distinct envelope tracing amplitude over time, which captures these complex temporal dynamics. **B.** Spectrogram of vocalization-modulated noise presented to one A1 neuron. Stimuli were generated by applying vocalization envelopes to narrowband noise, capturing the complex temporal dynamics of natural sounds. For the two-band stimulus, a different envelope was applied to adjacent, non-overlapping spectral bands. Both noise streams were positioned in the responsive area of a frequency tuning curve (right). Thus vocalization-modulated noise enabled probing natural, nonlinear temporal processing while minimizing complexity of spectral features. **C.** Raster response the same neuron to repeated presentations of the vocalization-modulated noise stimulus (top), and peri-stimulus time histogram (PSTH) response averaged across repetitions (gray shading, bottom). The envelope of each noise stream is overlaid. Increased amplitude in stream 2 (blue) leads to a strong onset response that weakens after about 50 ms (transients in the PSTH at 0.25 s and 0.8 s). Stream 1 (orange) suppresses the PSTH, with no evidence for adaptation.

The dynamic vocalization-modulated noise often evoked reliable time-varying responses from A1 neurons, but the timecourse of this response varied substantially. Peri-stimulus time histogram (PSTH) responses computed from average repetitions of identical noise stimuli showed that responses could predominantly follow the envelope of one or both of stimulus bands (Fig. 1C). Thus, while all neurons included in the study were excited by isolated, narrowband stimuli in each frequency band, responses to stimuli presented in both bands simultaneously were complex and varied across neurons.

We used a linear-nonlinear spectro-temporal receptive field model (LN STRF) to establish a baseline characterization of auditory encoding properties (Fig. 2A, [41–43]). This model describes time-varying neural activity as the linear weighted sum of the preceding stimulus spectrogram (Eq. 1). Because the vocalization-modulated noise consisted of just two distinct spectral channels, the STRF required a filter with only two input spectral channels, compared to multiple spectral channels for analysis of broadband noise or natural sounds. To account for well-established nonlinear threshold and saturation properties of spiking neurons, the linear filter stage was followed by a static, sigmoidal output nonlinearity (Eq. 2, [31], Fig. 2A).

**Figure 2.**
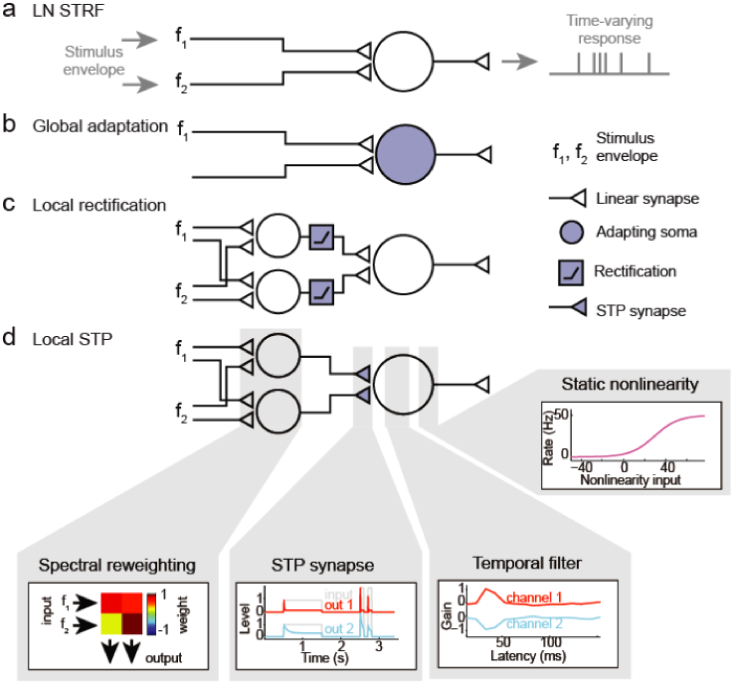
Alternative encoding models to describe auditory neural responses to vocalization-modulated noise. A. The linear-nonlinear spectro-temporal receptive field (LN STRF) describes the time-varying neural response as a linear weighted sum of the preceding stimulus envelopes, followed by a static sigmoid nonlinearity to account for spike threshold and saturation. B. In the global short-term plasticity (STP) STRF, nonlinear STP (depression or facilitation) is applied to the output of the linear filter prior to the static nonlinearity. C. In the local rectification STRF, the input channels are linearly reweighted and then nonlinearly thresholded (rectified) prior to the linear temporal filter and static nonlinearity. D. In the local STP STRF, input channels are linearly reweighted, and then nonlinear STP (depression or facilitation) is applied to each reweighted channel, prior to the linear temporal filter and static nonlinearity. Gray boxes show example model parameters applied at each processing stage.

**Figure 3.**
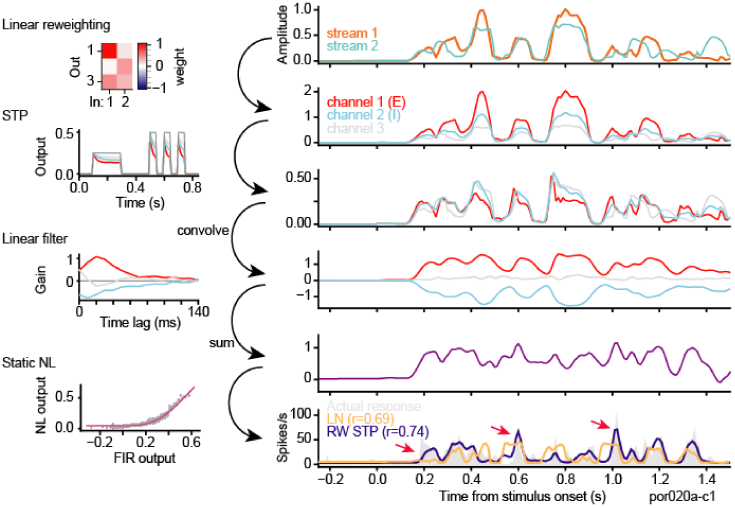
Transformation applied to incoming vocalization-modulated noise for a local short-term plasticity (STP) STRF estimated for one A1 neuron. Spectral reweighting emphasizes input stream 1 in channel 1 (red), stream 2 in channel 2 (blue), and both streams in channel 3 (gray). All three reweighted channels undergo independent STP. For this neuron, STP is stronger for channel 1 than for the other channels. The linear filter produces excitation for channel 1 and inhibition for channel 2. After the final static nonlinearity (NL), the predicted PSTH (bottom panel, purple) shows a good match to the actual PSTH (gray shading), while the prediction of the LN STRF does not predict the response dynamics as accurately (orange). Arrows indicate transient PSTH features captured better by the STP STRF.

To regularize model fits, we constrained the temporal dynamics of the filter applied to each input channel to have the form of a damped oscillator (Eq. 3, [31]). This parameterization required fewer free parameters than a simple, nonparametric weighting vector and improved model performance over a nonparametric linear filter (see Fig. 4C). However, the shape of the parameterized temporal filter did not capture temporal responses dynamics fully for all neurons. To support more flexible temporal encoding, we introduced a spectral reweighting in which the two input channels were mapped to *J* > 2 channels prior to temporal filtering (Eq. 4). Each reweighted input was passed through a separately-fit temporal filter. Several values of *J* were tested. For the majority of model comparisons, *J* = 5 was found to produce the best performing models on average and most results below are for models with this channel count (although *J* = 2 spectral channels achieved nearly asymptotic performance for the LN STRF, see Fig. 4C).

**Figure 4.**
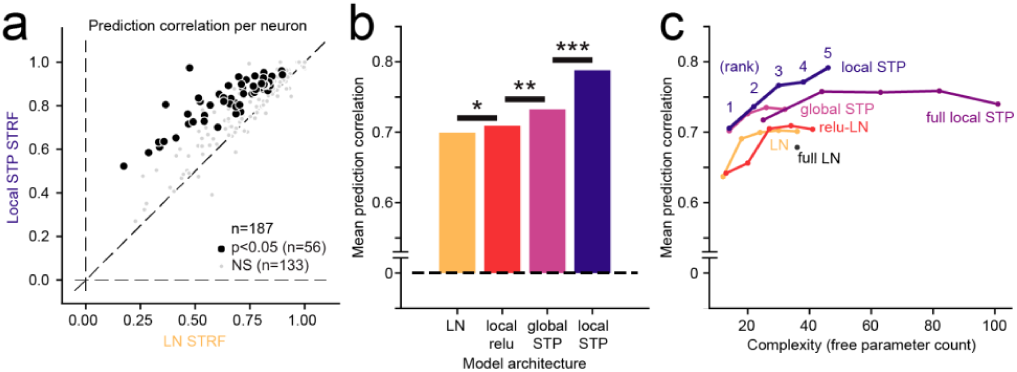
**A.** Scatter plot compares noise-corrected prediction correlation between the linear-nonlinear (LN) STRF and local short-term plasticity (STP) STRF for each A1 neuron. Black points indicate the 56/187 neurons for which the local STP model performed significantly better than the LN model (*p*<0.05, jackknifed *t*-test). **B.** Mean performance (noise-corrected correlation coefficient between predicted and actual PSTH) for each model across the set of A1 neurons. The global STP STRF showed improved performance over the LN and local rectification (relu) STRF. The local STP STRF showed a further improvement over the global STP STRF (\**p* < 0.01, \*\**p* < 10^-4^, \*\*\**p* < 10^-6^, Wilcoxon sign test, *n* = 187/200 neurons with above-chance prediction correlation for any model). The best performing model, the local STP STRF, reweighted the two input envelopes into five spectral channels, each of which underwent independent STP prior to linear temporal filtering and a static nonlinearity. **C.** Pareto plot compares model complexity (number of free parameters) versus average prediction correlation for model architectures with and without STP, with and without parameterization of the temporal filter (full vs. DO) and for variable numbers of reweighted spectral channels (rank). Models with STP showed (purple, blue) consistently better performance than models without STP (orange, red) for all levels of complexity.

The LN STRF, as well as the other models discussed below, was fit using gradient descent [44,45]. Model fits were regularized by the parametric formulation of the linear filter (Eqs. 3-4) and by a shrinkage term applied to the mean squared error cost function [31]. Model performance was assessed by the accuracy with which it predicted the time-varying response to a novel validation stimulus that was not used for estimation [4]. Prediction accuracy was quantified by the correlation coefficient (Pearson’s *R*) measured between the predicted and actual PSTH response, corrected to account for sampling limitations in the actual response [46]. A value of *R*=1 indicated a perfect prediction and *R*=0 indicated random prediction.

The LN STRF was able to capture some response dynamics of A1 neurons, but several errors in prediction can be seen in example data (Figs. 3, 5). In particular, the LN STRF failed to account for transient responses following stimulus onset (arrows in Fig. 3). A previous study showed that, for stimuli consisting of a single modulated noise band, a model incorporating nonlinear short-term synaptic plasticity (STP) prior to the temporal filtering stage provides a more accurate prediction of neural activity [17]. STP is widespread across cortical systems, making it a plausible mechanism to support such adaptation [17,26]. Given that STP occurs at synaptic inputs, this observation suggests that A1 neurons can undergo adaptation independently for inputs in different spectral channels. Spectrally tuned adaptation could give rise to a rich code for complex spectro-temporal patterns [29]. However, based on previous results, it is not clear whether the nonlinear adaptation occurs primarily after information is summed across spectral channels (global adaptation) or if it occurs separately for the different spectral channels (local adaptation). To determine whether adaptation occurs pre- or post-spectral integration, we estimated two variants of the STRF, a *global STP STRF*, in which input spectral channels undergo the same adaptation prior to linear filtering (Fig. 2B), and a *local STP STRF*, in which each channel adapts independently according to the history of its own input (Fig. 2D).

**Figure 5.**
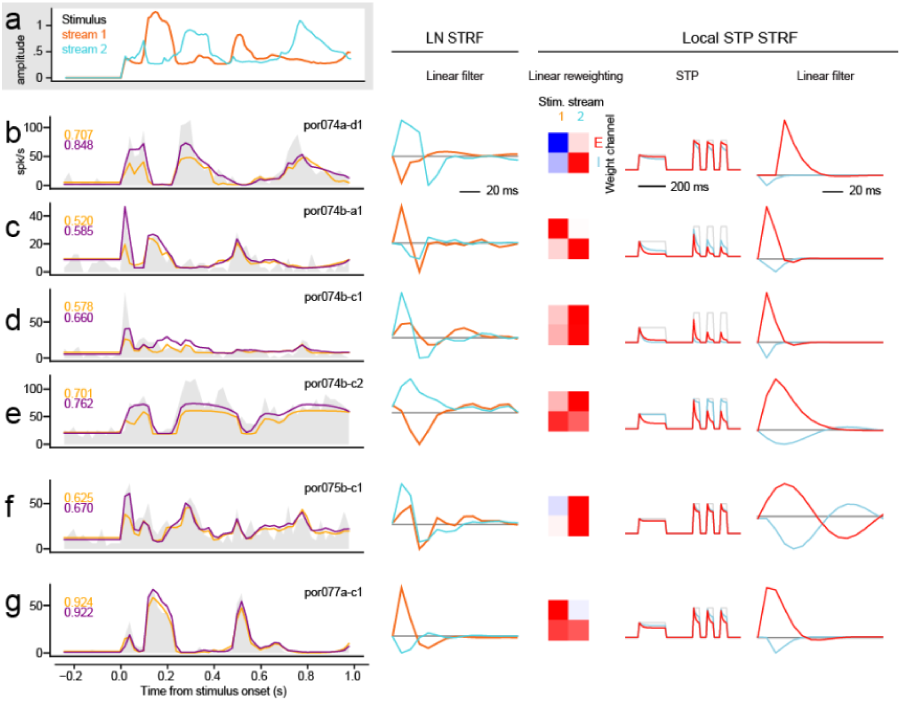
**A.** Envelope of vocalization-modulated noise streams. **B-G.** Left column, example PSTH responses of several A1 neurons (gray shading). The spectral position of noise bands was adjusted to fall within the receptive field of each neuron, but the envelopes were the same for each recording. Responses were sometimes dominated by one stream (e.g., unit C tracks stream 1 and G tracks stream 2), but could also track both (e.g., unit F). Response dynamics also vary substantially, from sustained, following the stimulus envelope (G), to highly transient responses that attenuate after sound onset (D). Numbers at upper left of PSTH plots indicate prediction correlation for the linear-nonlinear (LN) STRF (orange) and local short-term plasticity (STP) STRF (blue). Predicted PSTHs are overlaid on the actual PSTH. Second column shows linear filters from the LN STRF for each neuron, whose gain reciprocates the PSTH responses. Columns at right show spectral weights, STP properties and linear filters for the largest (positive gain, red) and smallest (negative gain, blue) temporal filter in local STP STRFs for the same neurons.

Spectral reweighting was applied to the stimulus for STP STRFs (Eq. 4), as in the case of the LN STRF, above. For the local STP STRF, nonlinear adaptation occurred after spectral reweighting. The reweighting made it possible for the same band of the vocalization-modulated noise to undergo adaptation at multiple timescales and, conversely, for different bands to be combined into a single channel before adaptation. This flexible arrangement models cortical neurons, where inputs from peripheral channels can be combined either pre- or post-synaptically [47,48]. The model schematic shows a model in which the two inputs were reweighted into two channels (Fig. 2D), but we compared models with *J* = 1…5 channels (see Fig. 4 and Methods). As in the case of the linear STRF, the STP models included the same sigmoidal output nonlinearity. The linear filter and static nonlinearity architectures were the same across LN and STP models, and all models were fit using identical data sets. However, the free parameters for each model were fit separately.

To test for the possibility that any benefit of the local STP STRF simply reflects the insertion of a nonlinearity into the LN STRF between spectral reweighting and temporal filtering, we considered an additional model, the local rectification STRF, in which each reweighted channel was linearly rectified prior to temporal filtering (Eq. 9, Fig. 2C).

These different encoding models can each be cast as a sequence of transformations, where the output of one transformation is the input to the next. Their modularity enables visualization of how the data is transformed at each step of the encoding process. Figure 3 illustrates the transformations that take place in an example local STP STRF for an A1 neuron (*J* = 3 spectral reweighting channels shown for simplicity). The vocalization-modulated noise envelope is first linearly reweighted into three channels. In this example, the first reweighted channel closely follows the first input channel. Second, the three reweighted channels undergo independent STP-like adaptation. The first channel experiences (red) the strongest adaptation. The adapted channels are then convolved with a linear filter, which in this case is excitatory for channel 1, inhibitory for channel 2 (blue), and transient excitation for channel 3 (gray). The convolved channels are summed and then passed through a static nonlinearity to generate the final predicted time-varying spike rate. The PSTH response predicted by the reweighted STP STRF can be compared directly to the actual PSTH and predictions by other models (Fig. 4).

For 187 out of the 200 A1 neurons studied, at least one STRF (LN, global STP, local rectification, local STP, *J* = 5 spectral reweighting channels for all models) was able to predict time-varying responses with greater than chance accuracy (*p*<0.05, Bonferroni-corrected permutation test). Prediction correlation for the global STP model was significantly greater than the linear model for a subset of neurons (*n* = 22/187, *p* < 0.05, permutation test, Fig. 4B). The average noise-corrected prediction correlation across the entire sample of neurons was greater for the global STP model (mean 0.699 vs. 0.732, median 0.715 vs. 0.755, *p* = 2.1 × 10^-7^, sign test). Mean performance tended to be slightly lower than median, probably because performance was near the upper bound of *r* = 1.0, creating a slight negative bias in the mean. However, we saw no qualitative difference between these metrics in any model comparison. The local rectification STRF also showed an average improvement in performance over the LN STRF (mean 0.699 vs. 0.709, median 0.715 vs. 0.732, *p* = 0.042, sign test, Fig. 4B). However, the local STP STRF consistently performed better than all the other models (mean 0.795, median 0.818, p<10^-8^ for all models, sign test, Fig. 4B). Prediction accuracy was significantly greater than the LN STRF for 58/187 neurons (*p* < 0.05, permutation test, Fig. 4A). Taken together, these results indicate that the spectrally tuned nonlinear adaptation described by the local STP STRF provides a more accurate characterization of A1 encoding than LN models or models in which the adaptation occurs uniformly across spectral channels.

While the local STP STRF consistently performed better than the other models, its performance could be attributed to its additional complexity, *i.e.*, the fact that it required more free parameters than the other models, rather than something specific about spectrally tuned adaptation. To characterize the interaction of model complexity and performance, we compared prediction accuracy for models with variable numbers of spectral reweighting channels, *J*=1…5 (Fig. 4C). When compared in a Pareto plot, the local STP STRF shows a consistent pattern of improved performance over LN STRFs, independent of spectral channel count or overall parameter count. This comparison also included models in which the temporal filter was either parameterized by a damped oscillator (Eq. 3) or nonparameterized (“full”). The parameterized models performed consistently as well or better than their nonparameterized counterparts, indicating that this reduction in dimensionality preserved important temporal filter properties. Thus, the benefit of incorporating local STP is consistent, regardless of model complexity.

### Spectrally tuned adaptation is stronger for excitatory than inhibitory inputs

We studied properties of the LN and STP STRFs in order to understand what features of the STP STRFs lead to their improved performance. Response dynamics varied across A1 neurons, sometimes emphasizing only sound onsets and in other cases tracking one or both envelopes across the entire trial. For many neurons, both models were able to capture the coarse response dynamics, but the STP model was able to predict the transient responses and the relative amplitude of responses more accurately (Fig. 5B-C). In some cases the LN and STP STRFs performed equivalently, indicating that some neurons showed little or no nonlinear adaptation (Fig. 5G).

Although isolated stimuli in both input channels usually evoked excitatory responses (Fig. 1C), the gain of one filter in both LN and STP STRFs was often negative (Fig. 5, middle and right columns). These suppressive responses likely reflect the unmasking of inhibition by broadband stimuli [49]. The fit procedure was not constrained to require a negative channel, so the presence of negative channels is the result of optimizing model parameters for prediction accuracy. We quantified the gain of each local STP STRF channel by summing temporal filter coefficients across time lags. By definition, one channel always had the largest gain, which we identified as the strongest input channel. A comparison of gain for largest versus smallest gain showed that one channel was always positive (*n* = 187/187 units, Fig. 6B). Strikingly, almost every filter contained at least one channel with negative gain (*n* = 175/187). We focus on models with *J*=5 spectral reweighting channels here, but nearly the same results are observed for models with *J* = 2…5. There was no difference in the prevalence of inhibitory channels in neurons that showed a significant improvement for the STP model, compared to neurons that did not show an improvement (*p* > 0.05, unpaired *t*-test). Although the specific mechanisms producing positive and negative gain are not determined in this model, we refer to them as excitatory and inhibitory gain, respectively.

**Figure 6.**
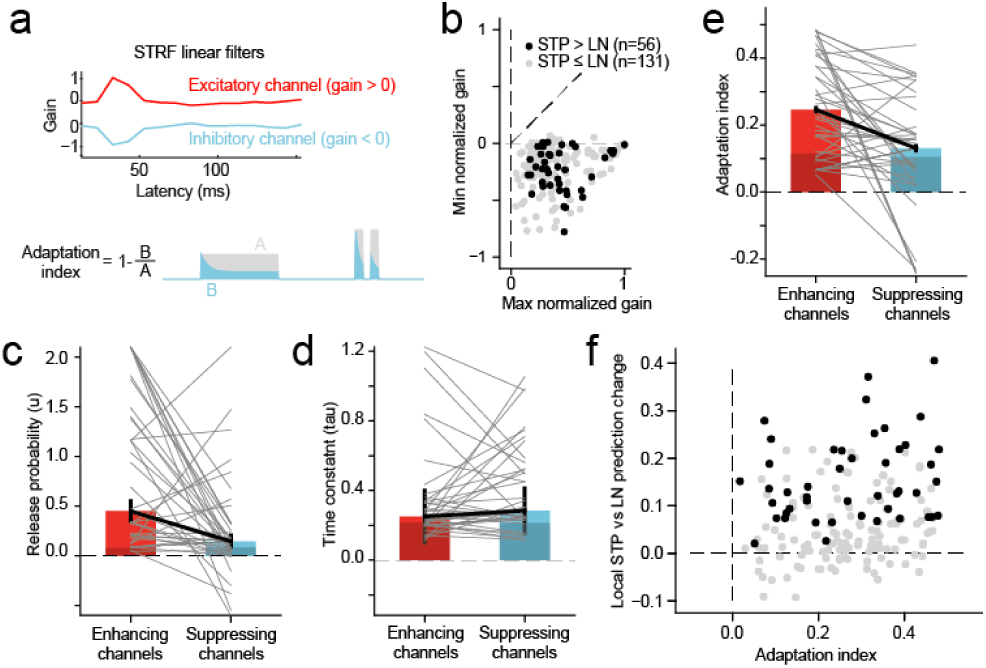
**A.** In the local STP STRF, each reweighted spectral channel passed through a nonlinear filter, mimicking synaptic STP, a nonlinear transformation, prior to the linear temporal filter stage. An index of adaptation strength for each model synapse was computed as one minus the fraction change in the amplitude of a test signal after passing through the adapting synapse. An index value > 0 indicated synaptic depression, and a value < 0 indicated facilitation. **B.** Overall gain for each channel of the linear filter in the STP STRF was computed as the sum of the filter across time lags. Scatter plot compares gain for the channel with largest magnitude, which was always positive (horizontal axis), and for the channel with smallest magnitude, which was either positive or negative (vertical axis). The vast majority of STRFs contained at least one excitatory (positive) channel and one inhibitory (negative) channel (n = 183/187). Units in which the local STP STRF generated a significant improvement in prediction power are colored black (n = 56, *p*<0.05, jackknife *t*-test). **C.** Comparison of release probability parameter fit values for STP filters in excitatory versus inhibitory channels (*n* = 56 STP STRFs with significant improvement in prediction power). Gray lines connect values for a single STRF. Average values were significantly greater for excitatory versus inhibitory synapses for release probability (mean 0.45 vs. 0.15, p = 1.4 × 10^-6^, sign test). **D.** Comparison of STP recovery time constant, plotted as in D, shows no difference between excitatory and inhibitory channels (mean 0.063 vs. 0.081 s, *p* > 0.5, sign test). **E.** Comparison of adaptation index shows a significant difference between excitatory and inhibitory channels (mean 0.25 vs. 0.13, *p* = 2.8 × 10^-4^ sign test). **F.** Scatter plot compares average adaptation index for each local STP STRF against the change in prediction correlation between the LN and local STP STRF. There is a positive correlation between STP effects and changes in prediction accuracy (*r* = 0.17, *p* = 0.023, n = 187, Wald Test). Neurons with significant changes in prediction accuracy are plotted as in B.

We wondered whether adaptation captured by the STP model differed between excitatory and inhibitory channels. For the 56 neurons with improved performance by the local STP STRF (see Fig. 4, above), we compared STP parameters (release probability and recovery time constant, see Eq. 6) and the overall adaptation index between highest- and lowest gain channels. The adaptation index was measured as one minus the ratio of the output to input of the synapse for a standard test input (Fig. 6A, [17]). Index values greater than zero indicated depression, and values less than zero indicated facilitation. When we compared STP properties between channels, we observed that release probability and adaptation index were both stronger, on average, for excitatory versus inhibitory channels (*p* = 0.0011 and *p* = 4.3 × 10^-4^, respectively, sign test, Fig. 6C, E). The mean adaptation index of excitatory channels (0.27) was more than twice that of inhibitory channels (0.13). These results suggest that excitatory responses in A1 tend to adapt following sustained input, while concurrent inhibition undergoes little or no adaptation. Mean recovery time constant did not differ between excitatory and inhibitory channels, possibly because the value of the time constant has little impact on model behavior when adaptation is weak (Fig. 6D).

We also tested whether the magnitude of STP-like adaptation predicted the relative performance of the local STP STRF. A comparison of average adaptation index versus change in prediction accuracy between the LN and local STP STRF for each neuron shows a small but significant correlation *(r* = 0.17, *p* = 0.023, *n* = 187, Wald Test for non-zero slope, Fig. 6F). When we considered neurons for which local STP STRF performance was not greater than the LN STRF, no mean difference was observed between excitatory and inhibitory channels (Fig. 6C-E, dark bars). However, the local STP models did tend to show non-zero STP strength, even if there was no significant improvement in performance. While many neurons did not show a significant improvement in prediction accuracy for the local STP STRF, the vast majority showed a trend toward improvement (168/187, Fig. 4A). If more data were available, permitting more robust STRF estimates, the number of neurons showing significant STP effects could be larger.

### Spectrally tuned adaptation supports contextual effects of stimulus-specific adaptation

Nonlinear adaptation has previously been proposed to play a role in contextual effects on auditory cortical responses [7]. One common measure of contextual influences on auditory activity is stimulus specific adaptation (SSA, [10,50]). When two discrete stimuli are presented in a regular sequence, with a standard stimulus presented more frequently than an oddball stimulus, responses to the standard tend to undergo adaptation, but responses to the oddball stimulus can be less adapted or even facilitated relative to a silent context. Effects of SSA have been attributed to feedforward adaptation and/or lateral inhibition [11–13,28].

To test for SSA effects, for a subset of neurons we presented standard/oddball sequences of noise bursts, falling in the same spectral bands as the vocalization-modulated noise stimuli. We measured SSA for these responses by an SSA index (SI) that compared responses to noise bursts when they appeared as standards vs. oddballs [10]. Adaptation effects were weaker than previously been reported for A1 in anesthetized animals, but SI was significantly greater than zero in 43% of neurons (*p*<0.05, standard/oddball permutation test, *n* = 44/102). We tested whether models including STP could predict responses to oddball stimuli and explain SSA effects. LN, global STP, and local STP STRFs were fit to data collected during the presentation of the oddball sequences. Because the design of the oddball stimulus experiments did not include repetitions of the same sequences, models were fit and tested using single trials. This design precluded correcting the prediction correlation for variability in the neural response [31,46], leading to comparatively lower correlation values than for the other stimulus sets. Nonetheless, the introduction of nonlinear model elements improved prediction accuracy. Between the LN STRF vs. local STP STRF, 46% of the cells responses were significantly better predicted by the local STP model (*p <* 0.05, jackknifed *t-*test, *n* = 47/102, Fig. 7A). Each model showed a significant improvement in accuracy over the simpler one (LN vs global STP STRF, *p* = 1.7 × 10^-4^; global STP STRF vs local STP STRF, *p* = 1.9 × 10^-13^, sign test, Fig. 7C). This pattern of improvement closely parallels the vocalization-modulated noise data (Fig. 4).

**Figure 7.**
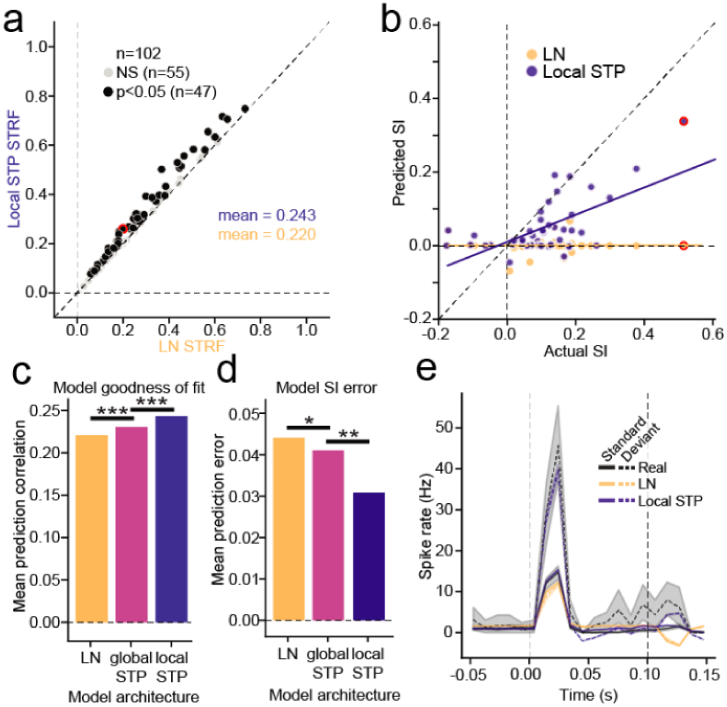
**A.** Scatter plot compares prediction accuracy for the LN STRF and local STP STRF, estimated using oddball stimuli for each neuron. Black markers indicate significant difference in the performance between models (*p*<0.05, jackknifed *t*-test). **B.** Scatter plot compares SSA index (SI) calculated from actual responses against SI from responses predicted by LN STRF (orange) and local STP STRF (blue) for neurons with significant actual SI (*p* < 0.05, standard/oddball permutation test). The LN STRF is unable to account for any stimulus specific adaptation, while the SI predicted by the local STP STRF is correlated with the actual values (LN: *r* = 0.011, *p* = 0.95; local STP: *r* = 0.636, *p* = 3.4 × 10^-6^, Wald Test for non-zero slope). **C.** Summary of the mean prediction correlation for all cells across all tested models (LN STRF vs. global STP STRF, *p* = 1.7 × 10^-4^, global STP STRF vs. local STP STRF, *p* = 1.9 × 10^-13^, LN STRF vs local STP STRF, *p* = 1.1 × 10^-15^, sign test). **D.** Mean SI prediction error for each model architecture. The prediction error for each cell is the mean standard error (MSE) between actual and predicted SI (LN STRF vs. global STP STRF, *p* = 0.024; global STP STRF vs. local STP STRF, *p* = 0.005, LN STRF vs. local STP STRF, *p* = 1.5 × 10^-4^, sign test). **E.** Example actual (black), LN STRF-predicted (yellow) and local STP STRF-predicted (blue) PSTH response to standard (continuous line) and deviant (dashed line) noise bursts. Shaded areas standard error on the mean (bootstrap *p* = 0.05). Vertical lines mark sound onset and offset. For the LN STRF, both standard and oddball predictions are close to the actual standard response, but the local STP STRF predicts the enhanced oddball response. Example cell is highlighted in panel A and B. \**p* <0 .05, \*\**p* < 0.01, \*\*\**p* < 0.001.

Because the models could predict time-varying responses to the noise stimuli, we could measure SI from responses predicted by the models. For neurons with significant SI (*p* < 0.05, permutation test, *n* = 44/102) we measured the correlation between actual and predicted SI values. The best performing model, the local STP STRF, was able to significantly predict the SI (*n* = 44, *r* = 0.636, *p* = 3.4 × 10^-6^, Wald Test for non-zero slope, Fig. 7B, blue). On the other hand, the LN STRF was unable to predict SI (*n* = 44, *r* = 0.011, *p* = 0.95, Wald Test, Fig 7B, orange). When comparing the SI prediction error across model architectures, the mean population error consistently decreased with the addition of spectrally tuned adaptation (LN vs global STP STRF, *p* = 0.024; global STP STRF vs local STP STRF, *p* = 0.005, sign test, Fig. 7D). Thus, A1 neurons that showed evidence for nonlinear STP-like adaptation also exhibited SSA, indicating that the two phenomena may share common mechanisms.

### Nonlinear adaptation is robust to changes in behavioral state

Several previous studies have shown that the response properties of neurons in A1 can be affected by changes in behavioral state. When animals engage in a task that require discrimination between sound categories, neurons can shift their gain and selectivity to enhance discriminability between the task-relevant categories [32,33,36]. Changes in overall gain are observed most commonly. Effects on sensory selectivity have been more variable and difficult to characterize.

We tested if changes in behavioral state influence the nonlinear STP-like adaptation we observed in A1. We trained ferrets to perform a tone detection task, in which they reported the occurrence of a pure tone target embedded in a vocalization-modulated noise sequence (Fig. 8A). We recorded neural activity during passive listening to the task stimuli and during active performance of the tone detection task. We then estimated STP STRFs in which the model parameters were either fixed between behavior conditions (passive listening versus active behavior) or allowed to vary between conditions. Because identical stimuli were used in both conditions, differences in the STRF could be attributed to changes in behavioral state. As in the case of SSA data, noise stimuli were not repeated within a behavioral block. Thus prediction accuracy was assessed with single trial data, and absolute prediction measures were lower than for the passive data reported above (Fig. 4).

**Figure 8.**
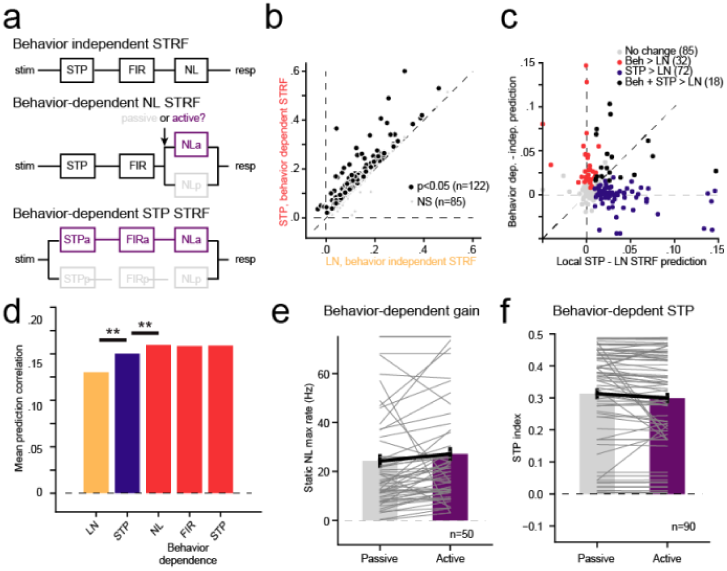
**A.** Schematic of alternative behavior-dependent local STP STRF models to account for changes in sound encoding between passive and active tone detection conditions. The behavior-independent model was fit independent of behavior state. For the behavior-dependent NL model, the static nonlinearity was fit separately for passive and active conditions but all other parameters were constant. Subsequent models introduced the active/passive split prior to earlier stages. **B.** Scatter plot compares prediction accuracy between the behavior-independent LN STRF and full behavior-dependent STP STRF for each cell in the set (pooled across on BF and away from BF target blocks). 122/207 neurons show a significant increase in prediction accuracy for the behavior-dependent model (*p*<0.05, jackknifed *t*-test). **C.** Relative change in prediction accuracy for each neuron from incorporating STP (LN vs. local STP STRF, x axis) versus incorporating behavior dependence (behavior independent vs. –dependent, y axis). The small number of units that show improvement for both models (black), is in the range expected by chance if STP and behavior effects are distributed independently across the A1 population (*p*>0.2, permutation test). **D.** Comparison of mean prediction accuracy for each model reveals a significant increase in performance for STP STRF over the LN STRF, as in the passive-only dataset in Fig. 4 (mean 0.13 vs. 0.15, *p* < 10^-10^). In addition, for the STP STRF, the behavior-dependent NL model shows improved performance over the behavior-independent model (mean 0.150 vs. 0.159, *p* = 2.2 × 10^-7^, sign test). However, no further improvement is observed if the linear filter or STP parameters are made behavior-dependent (*p*>0.05, sign test). **E.** Comparison of passive vs. active STRF gain (amplitude of the static nonlinearity) between active and passive conditions shows an increase in the mean response during behavior (mean NL amplitude 24 vs. 27 spikes/sec, *p* = 2.0 × 10^-5^, sign test). Gray lines show passive vs. active amplitude for each neuron. **F.** Comparison STP index for behavior-dependent STRF shows a small decrease in STP in the activity condition (mean 0.31 vs. 0.30, *p* = 0.002). This small behavior-dependent change does not impact mean prediction accuracy, as plotted in D.

When the parameters of the static nonlinearity were allowed to vary between passive and active states, allowing changes in gain between the passive and activate conditions, the models showed a significant improvement in predictive power when compared to the behavior-independent STRF (mean single trial prediction correlation 0.13 vs. 0.15, *p* = 7.1 × 10^-13^, sign test, Fig. 8B-D). However, allowing other model parameters to vary with behavioral state provided no additional improvement in model performance (*p* > 0.05, sign test, Fig. 8D). Thus, the changes in behavioral state influence the overall gain of the neural response without affecting the linear filter or nonlinear adaptation captured by the STP STRF.

We also considered whether the presence of STP-like adaptation in a neuron predicted its tendency to show behavior-dependent changes in activity. When we compared the incremental change in prediction accuracy resulting from addition of nonlinear STP or behavior-dependent gain to the STRF model, the relationship was highly variable (Fig. 8C). Some neurons showed improvement only for STP or behavior-dependence, and just a small number showed improvements for both. Overall, these effects occurred independently across the population (*p* > 0.1, permutation test). Thus the improved performance of the STP STRF does not predict the occurrence of behavior-dependent changes in activity.

The comparison of prediction accuracy between behavior-dependent models suggests that the response gain can change between passive and active conditions but STP parameters do not. When we compared parameters between STRFs fit separately under the different behavioral conditions, we found this to be largely the case. The average gain of the auditory response increased when animals engaged in behavior (mean amplitude of static nonlinearity: 24 vs. 27 spk/sec, *p* = 2 × 10^-5^, *n* = 50 neurons with significant improvement in behavior-dependent vs. behavior-independent STRF, sign test, Fig 8E). The average STP index showed a small decrease during task engagement (mean STP index 0.31 vs. 0.30, *p* = 0.0016, *n* = 10 neurons with significant improvement in local STP vs. LN STRF, sign test, Fig. 8F, right panels). While this change in STP index was significant, allowing it to fluctuate did not significantly impact prediction accuracy (Fig. 8D). A larger dataset may uncover significant influences of behavior-dependent nonlinear adaptation. However, the current analysis suggests that changes in STP play an overall smaller role in mediating behavioral effects in A1 than changes in overall response gain (Fig. 8E).

### Natural stimuli reveal nonlinear adaptation of spectrally overlapping channels in A1

The vocalization-modulated noise data reveal that spectrally distinct inputs can undergo independent adaptation in A1, supporting contextual coding phenomena such as SSA. In order to understand these nonlinear adaptation effects in a more ethological context, we also recorded the activity of 499 A1 neurons from 5 awake, passive ferrets during presentation of fully natural sounds. The natural stimuli were drawn from a large library of natural sounds (textures, ferret vocalizations, human speech and recordings of the ambient laboratory environment), chosen to sample a diverse range of spectro-temporal modulation space.

Neural encoding properties were modeled by a reduced-rank STRF model (Fig. 9A, [31,51]). For the LN STRF, the sound spectrogram passed through a bank of *J* = 4 spectral filters, each of which computed a linear weighted sum of the spectrogram at each time bin. The spectral filter output then passed through a linear temporal filter (constrained to be a damped oscillator, Eq. 3) and static nonlinearity, identical to elements in the vocalization-modulated noise models. To test for nonlinear adaptation, local STP was introduced to the model following the spectral filtering stage (Fig. 9A).

**Figure 9.**
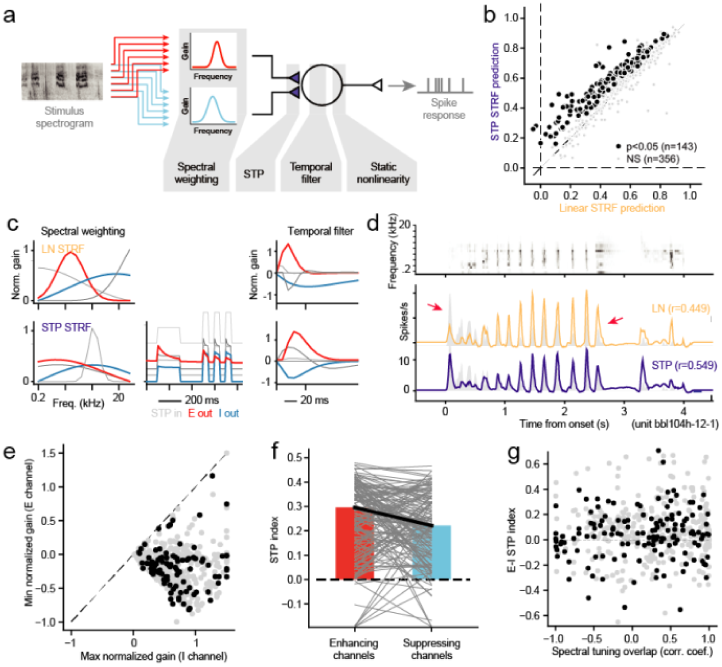
Performance of a local STP model on for A1 encoding of natural sounds. **A.** The encoding model for natural sounds resembled reweighted STP STRF for vocalization-modulated noise, except that the spectral filters at the first stage were two independently fit Gaussian functions that required two free parameters each (mean, standard deviation) and provided a simple tuning function for each spectral channel. **B.** Scatter plot compares prediction accuracy between the LN STRF and local STP STRF for the natural sound data. Across the entire set, 143/499 neurons showed a significant improvement in prediction accuracy for the local STP STRF (*p* < 0.05, jackknife *t*-test). Mean prediction accuracy for the local STP STRF was significantly greater than the LN STRF (0.517 vs. 0.563, *p* < 10^-10^, sign test). **C.** Example spectral weights and temporal filters for one LN STRF (top) and spectral weights, STP measure, and temporal filters for the local STP STRF for the same neuron. Maximum gain is normalized to 1, but the relative gain between channels is preserved. As is typical in the vocalization-modulated noise data, the highest gain filter (red) shows relatively strong STP, and the lowest gain (blue) shows weaker STP. **D.** Predicted PSTH responses for each model for one natural sound stimulus, overlaid on the actual PSTH (gray). The LN STRF prediction (orange) undershoots the initial transient response and over-predicts the sequence of transient responses later in the stimulus (arrows), while the STP STRF predicts these features more accurately (blue). **E.** Comparison of gain for the most positive (max normalized gain) and most negative (min normalized gain) linear filter channels for STP STRFs reveals that the majority fits contain one excitatory and one inhibitory channel. **F.** Comparison of STP strength between excitatory and inhibitory channels shows consistently stronger depression for the excitatory channels (mean 0.30 vs. 0.22, *p* = 4.1 × 10^-3^, sign test, *n* = 143 units with significant improvement for the STP STRF). **G.** Scatter plot compares overlap of E and I spectral channels for each STP STRF (x axis) and relative difference in STP index between the E and I channels. There is no correlation between tuning overlap and STP index difference, suggesting that A1 neurons represent incoming sound with a diverse combination of spectral tuning and nonlinear adaptation.

The STP STRF predicted time-varying natural sound responses more accurately, on average, than the LN STRF (Fig. 9B). The STP model performed significantly better for 143/499 of the A1 neurons studied, and the average prediction accuracy was significantly higher for the STP model (mean noise-corrected prediction correlation 0.517 vs. 0.563, median: 0.540 vs. 0.583, *p* < 10^-20^, sign test). Thus, introducing local nonlinear adaptation to a spectro-temporal model for encoding of natural sounds provides a similar benefit as for encoding of vocalization-modulated noise.

An example comparing LN and local STP STRF fits for one neuron shows a similar pattern of spectrally tuned adaptation as observed for the vocalization-modulated noise data (Fig. 9C). In this example, the spectral channel with strongest positive gain (red) shows relatively strong STP, while the channel with strongest negative gain (blue) shows very little evidence for STP. The net effects of this tuned STP can be observed in the predicted PSTH response to a natural sound (Fig. 9D). The LN STRF fails to predict the strong transient response at the sound onset and over-predicts the sequence of transients 1-2 sec after sound onset. The local STP STRF captures these dynamics more accurately.

As in the case of the vocalization-modulated noise data, we compared STP effects between excitatory and inhibitory channels. Temporal filters were ordered by their average gain, and the highest- and lowest gain filters were selected for comparison of STP properties (Fig. 9E). This comparison revealed that mean STP index was significantly larger for excitatory channels (mean 0.30) than for inhibitory channels (mean 0.22, *p* = 4.1 × 10^-3^, sign test, Fig. 9F). As in the case of vocalization-modulated noise (Fig. 6), the weaker STP for inhibitory channels suggests that these inputs tend to undergo little or no adaptation, while excitatory inputs undergo stronger adaptation. These effects did not depend on spectral tuning of the filters, as the differences in STP for excitatory versus inhibitory channels were consistent across filter center frequencies. There was also substantial heterogeneity in the strength of STP and the degree of overlap between spectral filters in an STRF (Fig. 9G). Thus, while many A1 neurons showed evidence for STP-like adaptation, especially in excitatory channels, the amount of adaptation and spectral overlap varied widely between neurons.

## Discussion

We found that the adaptation of neurons in primary auditory cortex (A1) to natural and naturalistic sounds is spectrally selective. These adaptation effects can be modeled by a neuron with multiple input synapses that independently undergo short-term plasticity (STP). Spectro-temporal receptive field (STRF) models that incorporate nonlinear, spectrally tuned adaptation predict neural responses more accurately than the classic linear-nonlinear (LN) STRF for both naturalistic vocalization-modulated noise and for fully natural stimuli. They also predict responses more accurately than models that undergo a global (non-spectrally tuned) adaptation. These adaptation effects are stable across changes in behavioral state, even as neurons undergo task-related changes in the gain of sound-evoked responses [32,34]. While the observed adaptation could be produced by a mechanism other than STP, these results demonstrate a general principle, that spectrally tuned adaptation plays an important role in encoding of complex sound features in auditory cortex. Across a variety of stimulus conditions (Figs. 4, 7, 9) and models of varying complexity (Figs. 4, 10), a simple STP-like mechanism provides a consistent improvement in the performance of encoding models.

**Figure 10.**
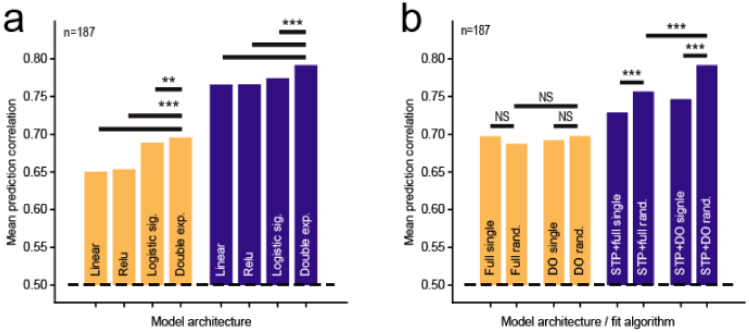
**A.** Impact of STRF output nonlinearity on prediction accuracy. Groups of bars compares mean prediction accuracy of LN (orange) and local STP STRFs (blue) with different output nonlinearities using the vocalization-modulated noise data. In both the LN and STP architectures, the double exponential sigmoid shows better performance than a model with no output nonlinearity (linear), linear rectification (relu), and a logistic sigmoid (\*\*\**p*<10^-5^; NS: *p*>0.05 sign test). **B.** Comparison of initialization method and parameterization on LN (orange) and local STP STRF (blue) performance. Full models used non-parameterized temporal filter functions, and DO indicates model in which the temporal filter is constrained to be a damped oscillator. Single fits started from a single initial condition, and random fits stated a 10 different initial conditions, selecting the best-performing model on the estimation data. Initialization and parameterization had little impact on LN STRF performance, but both random initialization and DO parameterization improved performance for the local STP STRF (\*\**p*<10^-4^; \*\*\**p*<10^-5^; NS: *p*>0.05 sign test).

Spectrally tuned adaptation may support perception of complex sound features, such as phonemes in speech and vocalizations [1], and may be of particular importance for hearing in noisy environments [52–54]. Evoked activity in A1 rarely has latency longer than 50 ms, but adaptation lasting several tens to hundreds of milliseconds can modulate these short-latency responses. Neurons that undergo adaptation will change their effective spectro-temporal tuning while non-adapting neurons will not. By comparing responses of adapting and non-adapting neurons, a decoder can infer information about stimuli at longer latencies [17]. Thus adaptation can operate as an encoding buffer, integrating stimulus information over a longer time window than the latency of the evoked response.

### Neural coding of auditory context

Studies of contextual effects on auditory neural coding have shown that the spectro-temporal selectivity can change with statistical properties of sound stimuli, including temporal regularity [10,50], contrast [14,18], intensity [16,55,56], and noisy backgrounds [52,53]. These contextual effects are typically measured in the steady state: neural activity is characterized during discrete epochs in which the statistical properties defining context are held constant. The current results suggest that the same mechanisms that affect activity in the steady state also operate dynamically during the encoding of complex natural stimuli. Spectrally tuned adaptation supports a rich spectro-temporal code in which a continuously changing sensory context, reflecting the previous 100-1000 ms, modulates short-latency (0-100 ms) responses to continuous natural sounds [57].

The timecourse of STP-like adaptation occurs over tens to hundreds of milliseconds, consistent with the timecourse of adaptation in encoding models that incorporate contrast gain control [18]. These effects may also share dynamics with models in which local sensory context of synthetic tone stimuli modulates sound-evoked activity [20,58]. Previous studies of gain control and contextual modulation have suggested, variously, that either feed-forward adaptation (in cortex or midbrain) or recurrent cortical circuits could shape the encoding of spectro-temporal sound features [18–20,59]. In all cases, relatively slow changes in stimulus contrast or power around the neurons receptive field can influence sensory selectivity. Thus mechanisms other than STP may be able to support adaptation with similar dynamics. Further study is required to determine if these different models are functionally equivalent or how feedforward and feedback elements of the auditory network contribute to this dynamic coding.

An adaptation-based contextual code produced by mechanisms such as STP may extend broadly across the brain [29,60]. As a general computation, this nonlinear adaptation may serve to remove temporal correlations from upstream inputs. Theoretical studies of the visual cortex have argued that variation in synaptic depression across neurons can explain differences temporal frequency tuning across neurons [30,39]. Synaptic depression has also been implicated in producing gain control in hippocampus [60]. Thus an auditory code that uses spectrally tuned adaptation provides an example of a computational process that may occur generally across neural systems.

### Dynamic reweighting of excitatory and inhibitory input

While the STP model used in this study supported both depression and facilitation, the vast majority of measured adaptation effects were consistent with depression. Moreover, the strength of depression was generally much stronger for spectral inputs that produced an increase rather than decrease in neural firing rate. While the underlying mechanisms producing increases versus decreases in spike rate cannot be fully determined from extracellular recordings, we interpret these components of the STRF algorithmically as excitatory versus inhibitory responses, respectively. The predominance of adaptation in excitatory channels is consistent with a coding system in which responses to the onset of sound are broadly tuned, but as excitation adapts, the sustained inhibition sculpts responses so that sustained activity is tuned to a narrower set of sound features [61]. It has been established that the precise timing and relative strength of inhibition versus excitation can substantially impact tuning in A1 [62]; thus, dynamic changes in their relative strength during natural sound processing could substantially change encoding properties compared to what is measured in more traditional stimulus paradigms.

The relative tuning, strength, and adaptation properties of excitatory versus inhibitory inputs are not stereotyped, but instead they vary substantially across A1 neurons. In most neurons, the STP STRF revealed at least partially overlapping excitatory and inhibitory inputs (Fig. 9), consistent with previous work [62,63]. However, across individual neurons, the best frequency and bandwidth of excitatory channels can be greater or smaller than those of the inhibitory channels. Thus, instead of reflecting a fixed pattern of selectivity, neurons display a diversity of tuning properties that supports a rich code of distinct spectro-temporal patterns. This diversity of synaptic properties may explain the differences in selectivity across A1, including spectro-temporal tuning [64], monotonic versus non-monotonic level tuning [65,66] and temporal versus rate coding of temporal modulations [67,68].

The present study was performed on serial recordings of isolated single units, ignoring possible interactions between neurons that could influence sound coding [69]. Simultaneous recordings of neural populations will illuminate the role of adaptation on network connectivity and population dynamics that likely contribute to context-dependent encoding [70,71].

### Minimal complexity for auditory encoding models

A broad goal motivating this study is to identify the essential computational elements that support nonlinear sound encoding in auditory cortex, in particular, under natural stimulus conditions. While several complex, nonlinear models have been shown to predict auditory neural activity better than the LN STRF [18–25], no single model has been adopted widely as a new standard. One reason a replacement has not been identified may simply be that the auditory system is complex and that current findings are not exhaustive enough to determine a single model that generalizes across stimulus conditions, species, and behavioral states. Indeed, only a few encoding models have been tested with natural stimuli [22], and these tests have often been performed in anesthetized animals [19,21]. In addition, each proposed model is built around different nonlinear elements, but it is likely that they existing in overlapping functional domains. That is, two different models may both perform better than the STRF because they capture the same adaptation process or nonlinear scaling of response gain. A comprehensive comparison of models using the same natural sound data set will help determine the best performing models and their degree of equivalence. To support such an effort, data from this study is publicly available, and the open source toolbox used for model fitting has a modular design, allowing testing of other model architectures in the same computational framework [45].

The current study took additional steps for testing an encoding model that have not typically been taken in previous studies. First, the local STP model was tested using multiple different types of stimuli (vocalization-modulated noise, oddball sequences, natural sounds), and it was shown to perform better than the LN and global STP STRF across stimulus conditions. Second, it compared models of varying complexity. For both low- and high-parameter count models, the addition of a relatively simple STP component provides an improvement in performance. Previous models have suggested that nonlinear adaptation can improve encoding model performance [19,21]. These alternative models are quite high-dimensional compared to standard LN STRF formulations. This study supports the adaptation hypothesis, but it also shows that the adaptation can be implemented with a small number of additional free parameters, as long as it occurs independently for input spectral channels.

### Stimulus specific adaptation

Stimulus specific adaptation is one of the best-studied contextual effects in auditory cortex [10,12,13,50]. The STP STRF developed in the current study is able to account for SSA during steady state sound presentation. At the same time, the STP model reveals that the same adaptation mechanisms support a broader dynamic code, in which the degree of adaptation is continuously updated to reflect the history of the changing stimulus. This adaptation represents a generalization of SSA, as it does not depend strictly on the regularity the sensory input [72]. In this way, STP parameters provide a complementary metric to SSA, able to explain nonlinear adaptation for a broader set of stimuli and readily scalable to analysis at a population level.

While nonlinear adaptation and SSA effects are correlated, the strength of this relationship varies across individual neurons. This variability supports the possibility that mechanisms other than STP contribute to SSA. The idea that synaptic depression alone can support SSA has been also disputed because oddball stimuli can sometimes evoked responses that are enhanced relative to those stimuli presented in isolation [28]. However, for a neuron with inhibitory inputs that undergo adaptation, a recurrent disinhibition mechanism could produce enhanced oddball responses [11]. The data in the current study suggest that inhibitory inputs generally show weaker adaptation than their excitatory partners, which is consistent with other modeling studies [61]. However, even inhibitory inputs do tend to undergo some depression, leaving open the possibility that they could explain the enhanced oddball responses during SSA. Inhibitory interneurons in auditory cortex have been shown to contribute to SSA [11], but their role in natural sound coding has yet to be characterized.

### Robustness of adaptation effects across changes in behavioral state

Studies in behaving animals have shown that gain and selectivity of A1 neurons can be influenced by changes in behavioral state, such as arousal, task engagement, and selective attention [33–36]. We observed changes in response gain during task engagement, consistent with this previous work, and incorporating behavior state-dependent gain into the STRF model improved prediction accuracy. However, average adaptation properties did not change across behavioral conditions. Moreover, allowing nonlinear adaptation to vary between behavior conditions did not improve model performance. Thus, STP-like adaptation properties appear to be largely stable across top-down changes in behavioral state. It remains to be seen if they change over longer time scales, but the relative stability of tuning suggests that nonlinear adaptation contributes to a veridical code of sound features in A1 that is selectively gated into high-order, behavior-dependent features in downstream auditory fields [73].

The approach of incorporating behavioral state variables into sensory encoding models may be useful for integrating bottom-up and top-down coding more broadly [74]. As sound features take on different behavioral meanings, such as when selective attention is engaged, coding in the auditory system must also shift to represent the behaviorally relevant sound features [75,76]. A complete understanding of state-dependent changes in sound encoding thus requires models of how neurons change their coding properties in different behavioral states.

## Materials and Methods

All procedures were approved by the Oregon Health and Science University Institutional Animal Care and Use Committee and conform to standards of the Association for Assessment and Accreditation of Laboratory Animal Care (AAALAC).

### Animal preparation

Eleven young adult male and female ferrets were obtained from an animal supplier (Marshall Farms, New York). A sterile surgery was performed under isoflurane anesthesia to mount a post for subsequent head fixation and to expose a small portion of the skull for access to auditory cortex. The head post was surrounded by dental acrylic or Charisma composite, which bonded to the skull and to a set of stainless steel screws embedded in the skull. Following surgery, animals were treated with prophylactic antibiotics and analgesics under the supervision of University veterinary staff. The wound was cleaned and bandaged during a recovery period. Starting after recovery from implant surgery (about two weeks), each ferret was gradually acclimated to head fixation using a custom stereotaxic apparatus in a plexiglass tube. Habituation sessions initially lasted for 5 minutes and increased by increments of 5-10 minutes until the ferret lay comfortably for at least one hour.

### Acoustic stimulation

Five awake, passively listening, head-fixed animals were presented with vocalization-modulated noise [17,34,40] (Fig. 1). The stimuli consisted of two streams of narrowband noise (0.25-0.5 octave, 65 dB peak SPL, 3 s duration). Each stream was centered at a different frequency and modulated by a different envelope taken from one of 30 human speech recordings [77] or ferret vocalizations from a library of kit distress calls and adult play and aggression calls [17]. Envelopes were calculated by rectifying the raw sound waveform, smoothing and downsampling to 300 Hz. Each envelope fluctuated between 0 and 65 dB SPL, and its temporal modulation power spectrum emphasized low frequency modulations, with 30 dB attenuation at 10 Hz, typical of mammalian vocalizations [78]. Thus, the spectral properties of the noise streams were simple and sparse, while the temporal properties matched those of ethological natural sounds.

For three animals, vocalization-modulated noise was presented during passive listening and during active performance of a tone detection task (see below). During passive experiments, both noise streams were positioned in non-overlapping frequency bands in a neuron’s receptive field (0.25-1 octave center frequency separation) and were presented from a single spatial location, 30 deg contralateral from the recorded hemisphere. During behavioral experiments, the streams were centered at different frequencies (0.9-4.3 octave separation) and presented from different spatial locations (±30 degrees azimuth), such that one stream fell outside of the spectral tuning curve. Spectral properties of the individual vocalization-modulated noise streams were otherwise identical to those used in the passive experiments above.

In a subset of experiments (two animals), an oddball stimulus was presented to passively listening animals to characterize stimulus-specific adaptation [10]. Stimuli consisted of a sequence of regularly repeating noise bursts (100 ms duration, 30 Hz), with the same center frequency and bandwidth as the vocalization-modulated noise presented during the same experiment. On each 20-second trial, 90% of the noise bursts fell in one band (standard) and a random 10% were in the other band (oddball). The spectral bands of the standard and oddball streams were reversed randomly between trials.

Finally, in a different set of experiments, six passively listening animals were presented a library of 93, 3-sec natural sounds. The natural sounds included human speech, ferret and other species’ vocalizations, natural environmental sounds, and sounds from the animals’ laboratory environment.

In all experiments, the majority of stimuli (28 vocalization-modulated noise samples and 90 natural sounds) were presented a few times (2-5 repetitions). The remaining samples from each sound library (2 vocalization-modulated noise samples and 3 natural sounds) were presented 10-30 times, allowing for robust measurement of a peri-stimulus time histogram (PSTH) response (Fig. 2). These high-repeat stimuli were used for measuring model prediction accuracy (see below).

Experiments took place in a sound-attenuating chamber (Gretch-Ken) with a custom double-wall insert. Stimulus presentation and behavior were controlled by custom software (Matlab). Digital acoustic signals were transformed to analog (National Instruments), amplified (Crown D-75A), and delivered through free-field speakers (Manger W05, 50-35,000 Hz flat gain) positioned ±30 degrees azimuth and 80 cm distant from the animal. Sound level was calibrated against a standard reference (Brüel & Kjær 4191). Stimuli were presented with 10ms cos^2^ onset and offset ramps.

### Neurophysiological recording

After animals were prepared for experiments, we opened a small craniotomy over primary auditory cortex (A1). Extracellular neurophysiological activity was recorded using 1-4 independently positioned tungsten microelectrodes (FHC). Amplified (AM Systems) and digitized (National Instruments) signals were stored using MANTA open-source data acquisition software [79]. Recording sites were confirmed as being in A1 based on dorsal-ventral, high-to-low frequency tonotopy and relatively reliable and simple response properties [80,81]. Some units may have been recorded from AAF, particularly in the high frequency region where tonotopic maps converge. Single units were sorted offline by bandpass filtering the raw trace (300-6000 Hz) and then applying PCA-based clustering algorithm to spike-threshold events [43]. Neurons were considered isolated single units if standard deviation of spike amplitude was at least two times the noise floor, corresponding to > 95% isolation of spikes.

A pure-tone or broadband noise probe stimulus was played periodically to search for sound-activated neurons during electrode positioning. Upon unit isolation, a series of brief (100-ms duration, 100-ms interstimulus interval, 65 dB SPL) quarter-octave noise bursts was used to determine the range of frequencies that evoked a response and the best frequency (BF) that drove the strongest response. If a neuron did not respond to the noise bursts, the electrode was moved to a new recording depth. Thus our yield of 187/200 neurons responsive to vocalization-modulated noise overestimates the rate of responsiveness across the entire A1 population. Center frequencies of the vocalization-modulated noise stimuli were then selected based on this tuning curve, so that one or both of the noise bands fell in the frequency tuning curve measured with single noise bursts.

### Tone detection task

Three ferrets were trained to perform a tone in noise detection task [34]. The task used a go/no-go paradigm, in which animals were required to refrain from licking a water spout during presentation of vocalization-modulated noise until they heard the target tone (0.5 s duration, 0.1 s ramp) centered in one noise band at a random time (1, 1.5, 2, … or 5 s) after noise onset. To prevent timing strategies, the target time was distributed randomly with a flat hazard function [82]. Target times varied across presentations of the same noise distractors so that animals could not use features in the noise to predict target onset.

In a block of behavioral trials, the target tone matched the center frequency and spatial position of one noise stream. Behavioral performance was quantified by hit rate (correct responses to targets vs. misses), false alarm rate (incorrect responses prior to the target), and a discrimination index (DI) that measured the area under the receiver operating characteristic (ROC) curve for hits and false alarms [34,83]. A DI of 1.0 reflected perfect discriminability and 0.5 reflected chance performance. A detailed analysis of behavior is reported elsewhere [34]. In the current study, only data from blocks with DI significantly greater than chance and correct trials were included in the analysis of neural encoding.

During recordings, one noise stream was centered over a recorded neuron’s best frequency and the other was separated by 1-2 octaves. The target tone fell in only one stream on a single block of trials. Identical task stimuli were also presented during a passive condition, interleaved with behavioral blocks, during which period licking had no effect. Previous work compared activity between conditions when attention was directed into versus away from the neuron’s receptive field [34]. Because of the relatively small number of neurons showing both task-related and STP effects in the current study (see Fig. 8), data were collapsed across the different target conditions. Instead, neural activity was compared for the vocalization-modulated noise stimuli between active and passive listening conditions.

### Spectro-temporal receptive field models

#### Linear-nonlinear spectro-temporal receptive field (LN STRF)

Vocalization-modulated noise was designed so that the random fluctuations in the two spectral channels could be used to measure spectro-temporal encoding properties. The *LN STRF* is a widely viewed as a current standard model for early stages of auditory processing [42,84–86]. The LN STRF is an implementation of the generalized linear model (GLM), which is used widely across the auditory and other sensory systems [87,88]. In the first, linear stage of this model, a finite impulse response (FIR) filter, *h*(*x,u*), is applied to the stimulus spectrogram, *s*(*x,t*), to produce a linear prediction of time-varying spike rate, *r*_*L*_(*t*),

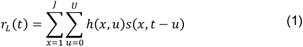

For the current study, the time lag of temporal integration, *u*, ranged from 0 to 150 ms. In typical STRF analysis, the stimulus is broadband and variable across multiple spectral channels, *x*. Here, the stimulus spectrogram was composed of just two time-varying channels, and a simplified version of the STRF was constructed in which *x* spanned just these two channels (*i.e., J* = 2), but we used larger values following spectral reweighting, below. A log compression was applied to the spectrogram to account for cochlear nonlinearities (offset 1 to force the compressed output to have nonnegative values, [31]). Otherwise, this model functions as a traditional STRF.

In the second stage of the LN model, the output of the linear filter, *r*_*L*_(*t*) is transformed by a static, sigmoidal nonlinearity, which accounts for spike threshold and saturation. The current study used a double exponential sigmoid,

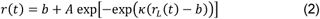

where *r*_0_ is the baseline (spontaneous) spike rate, *A* is the maximum evoked rate, *κ* is the slope, and *b* is the baseline. The specific formulation of the output nonlinearity does not substantially impact relative performance of models in which other aspects of model architecture are manipulated (see Output nonlinearity controls, below).

#### Temporal filter parameterization and spectral reweighting

As models become more complex (i.e., require fitting more free parameters), they become more susceptible estimation noise. While our fitting algorithm was designed to prevent overfitting to noise (see below), we found that constraining the temporal form of the linear filter in Eq. 1 improved performance over a model in which the filter was simple as set of weights for each time lag. Each spectral channel of the linear filter was constrained to be a damped oscillator, *h*_DO_(*x, u*),

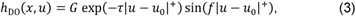

Requiring four free parameters: gain, *G*; latency, *u*_0_; duration, *tau*; and modulation frequency, *f*.

Because the damped oscillator constrains temporal tuning, we then considered the possibility that more than *J* = 2 spectral channels might be optimal for explaining neural responses to the two band vocalization-modulated noise stimulus. To allow for more than two spectral channels, we defined a reweighted stimulus, *s*_*R*_(*j,t*), computed as the input stimulus scaled by coefficients, *w*(*i,j*),

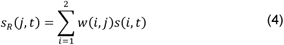

where *j* = 1…*J* maps the stimulus to a *J*-dimensional space. This reweighted stimulus provides input to the a damped oscillator, now also with J channels,

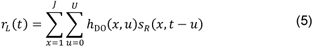

For the current study (except for controls, see below), the output of Eq. 5 is then transformed by the output nonlinearity (Eq. 2) to produce a predicted time-varying spike rate.

While we describe the LN STRF as a sequence of linear transformations — spectral filtering followed by temporal filtering—these two stages can be combined into a single linear spectro-temporal filter. We describe them as separate stages to frame the local STP model, below, where nonlinear adaptation is inserted between the two linear filtering stages.

#### Local short-term plasticity (STP) STRF

As several studies have demonstrated, the LN STRF captures import aspects of spectro-temporal coding but fails to account completely for time-varying sound evoked activity in auditory cortex [18,20,25,85]. In particular, the LN STRF fails to account for the temporal dynamics of sound-evoked activity [17,43]. Short-term synaptic plasticity (STP), the depression or facilitation of synaptic efficacy following repeated activation, has been proposed as one mechanism for nonlinear dynamics in neural networks [9,26]. A previous study showed that an LN model for A1 that incorporated STP was able to better explain the dynamics of responses to a single noise band with natural temporal modulations [17]. However, because that study utilized vocalization-modulated noise comprised of only a single noise band, it was not clear whether the nonlinear adaptation was *global*, affecting responses to all stimuli equally, or *local*, affecting only a subset of inputs independently. Because the current study used multiple noise channels, it could compare a global STP STRF, in which adaptation affected all input channels, to a local STP STRF, in which adaptation was spectrally tuned and could affect just a subset of inputs.

The effects of nonlinear adaptation were captured with a simple, two-parameter model of STP [26],

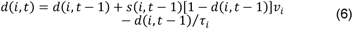

where *d*(*i,t*) describes the change in gain for stimulus channel *i* at time *t*. The change in available synaptic resources (release probability), *ν*_*i*_, captures the strength of plasticity, and the recovery time constant, *τ*_*i*_, determines how quickly the plasticity returns to baseline. Values of *d* < 1 correspond to depression (driven by *ν*_*i*_ > 0) and *d* > 1 correspond to facilitation (*ν*_*i*_ < 0). In the *local STP STRF*, each input channel of the stimulus is scaled by *d*(*i,t*) computed for that channel,

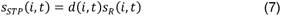

This nonlinearly filtered stimulus is then provided as input to the LN filter (Eqs. 5, 2) to predict the time-varying response. Note that if the strength of STP is 0 (i.e.,), then the STP STRF reduces to the LN STRF.

We also note that the local STP model uses the reweighted stimulus as its input. The reweighting allows the model to account for adaptation at multiple timescales on inputs from the same spectral band. Although the input is comprised of just two channels, the subsequent nonlinear filtering means that allowing the reweighted stimulus channel count, *J*, or rank, to be greater than two can increase model predictive power. In the current study, we evaluated models with rank *J* = 1-5. Predictive power was highest for *J* = 5. Higher values of *J* could, in theory, produce even better performance, but we did not observe further improvements for the current dataset.

#### Global STP STRF

We considered two control models to test for the specific benefit of spectrally tuned adaptation on model performance. One possible alternative is that a single, global adaptation is able to account for nonlinear temporal dynamics. To model global adaptation, the *global STP STRF* applied STP to the output of the linear filter (Eq. 5) before applying the static nonlinearity (Eq. 2). Thus, a single adaptation term was applied to all incoming stimuli, rather than allowing for the channel-specific adaptation in the local STP STRF. There is no simple biophysical interpretation of the global STP mechanism, but it can be thought of as a postsynaptic effect, capturing nonlinear dynamics similar to STP, but after integration across spectral channels. We compared performance of this model to a variant in which stimulus gain is averaged across spectral channels before scaling the input stimulus,

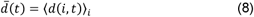

We found no difference between this common input STP model and the global STP STRF. Because the global model required fewer free parameters, we focused on this model for the comparisons in this study.

#### Local rectification STRF

Although spectral reweighting can improve the performance of the LN STRF by increasing the rank of the linear filter, it can still only account for linear transformations of the input stimulus. The STP model could, in theory, benefit simply from the fact that reweighted spectral inputs undergo any nonlinear transformation prior to the temporal filter. To control for the possibility that the STP nonlinearity is not specifically beneficial to model performance, we developed a local rectification STRF, in which the reweighted spectral inputs were linearly rectified with threshold *s*_0_ prior to temporal filtering,

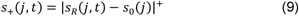

The rectified reweighted stimulus then provided the input to the LN model specified in Eqs. 7 and 2.

The set of encoding models described above represents a hierarchy of model architectures with increasing complexity, in that each successive model requires additional free parameters. Each model can be cast as a sequence of transformations applied to the stimulus, and the output of the final transformation is the predicted time-varying response (Fig. 3).

#### Fit procedure

Spike rate data and stimulus spectrograms were binned at 10 ms before analysis (no smoothing). The entire parameter set was fit separately for each model architecture. Data preprocessing, model fitting, and model validation were performed using the NEMS library in Python [45]. Identical estimation data from each neuron and the same gradient descent algorithm were used for each model (L-BFGS-B, [44]). The optimization minimized mean squared error (MSE) with shrinkage, a form of early stopping in which the standard MSE value is scaled by its standard error [31]. The use of parameterized temporal filters provided an effective regularization, as it constrained the shape of the temporal filter to be smooth and sinusoidal. Models were initialized at 10 random initial conditions (except for local minimum controls, see below), and the final model was selected as the one that produced the lowest MSE with shrinkage for the estimation data.

The ability of the encoding model to describe a neuron’s function was assessed by measuring the accuracy with which it predicted time varying activity in a held-out validation dataset that was not used for model estimation. The *prediction correlation* was computed as the correlation coefficient (Pearson’s *R*) between the predicted and actual PSTH response. Raw correlation scores were corrected to account for sampling limitations that produce noise in the actual response [46]. A prediction correlation of *R*=1 indicated perfect prediction accuracy, and a value of *R*=0 indicated chance performance. All models were fit and tested using the same estimation and validation data sets. Significant differences in prediction accuracy across the neural population were determined by a Wilcoxon sign test.

In a previous study involving just a single stream of vocalization-modulated noise, we tested our fitting procedure on simulated data produced by either an LN STRF or STP STRF. The estimated STRFs captured the presence or absence of the STP nonlinearity accurately [17]. In addition, the simulations revealed that LN STRFs could capture some aspects of the nonlinear adapting data, but estimated temporal filter properties did not match the actual temporal filter properties.

For data from the behavior experiments, which was all fit using single trials, 10-fold cross validation was used, on top of the procedure described above. Ten interleaved, non-overlapping validation subsets were drawn from the entire passive plus active data. The above fit algorithm was then applied to corresponding 90% estimation set, and the resulting STRF was used to predict the validation subset. Prediction accuracy was assessed for the conjunction of the 10 validation sets. Model parameters were largely consistent across estimation sets and average fit values are reported in the Results.

#### Output nonlinearity control

In a previous study, we compared performance of a variety of different static nonlinearities for LN STRFs and found that the double exponential sigmoid (Eq. 2) performed slightly, but consistently, better than other formulations of the output nonlinearity for A1 encoding models fit using natural vocalization stimuli [31]. We performed a similar comparison using the speech-modulated vocalization data, comparing LN and local STP STRFs with four different output functions (Fig. 10A): linear pass-through,

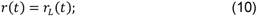

linear rectification,

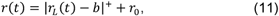

with threshold *b* and spontaneous rate *r*_0_; logistic sigmoid [18],

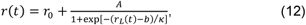

and the double exponential sigmoid (Eq. 2). As in the previous study, the double exponential sigmoid performed best for the LN model. The STP model incorporating a given output function always performed better than the corresponding LN model with the same output function, and the double exponential performed best overall. Thus for the rest of the study, we focused on models using the double exponential nonlinearity.

#### Temporal parameterization control

The use of a damped oscillator to constrain model temporal dynamics could, in theory, be suboptimal for describing neural response dynamics. We compared performance of the damped oscillator models to LN and local STP STRFs with nonparametric temporal filters, that is, where the temporal filter was simply a vector of weights convolved with the stimulus at each time point (Fig. 10B). For the LN STRF, the parameterization had no significant impact on average model performance (*p* > 0.05, sign test). For the local STP STRF, performance was higher for the parameterized model (*p* < 10-5, sign test), indicating that the parameterization was an effective form of regularization.

#### Local minimum control

While it has been shown that linear filters are well-behaved (*i.e.*, convex) and thus not subject to problems of local minima during fitting, it is more difficult to determine if local minima are adversely affecting performance of nonlinear or parametric models, such as the STP and damped oscillator, respectively, used in the current study. To determine if these models were negatively impacted by local minima during fitting, we compared performance of models fit from a single initial condition to the best model (determined using only estimation data) starting from 10 random initial conditions. Performance was compared for 4 different architectures: LN and local STP STRFs, each with parametric (damped oscillator) or nonparametric temporal filters (Fig. 10B). Each model was tested with the validation data. For the LN models, we saw no significant effect of using multiple initial conditions. For the local STP model, we saw a small but significant improvement when multiple initial conditions were used. Thus for the majority of results presented here, models were fit using 10 random initial conditions.

### Stimulus specific adaptation analysis

Sound-evoked activity recorded during presentation of the oddball noise burst sequences was modeled using the LN STRF, global STP STRF, and local STP STRF, as described above. To assess stimulus specific adaptation (SSA), an SSA index (SI) was used to measure the relative enhancement of responses to oddball versus standard noise bursts [10,50].

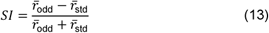

Here 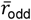 and 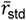 are the average response across bursts of both center frequencies, in the oddball and standard conditions, respectively. Neuronal response was calculated as the integral over time of the PSTH during the sound presentation. Significance SI was calculated with a shuffle test in which the identity of tones (oddball or standard) was randomly swapped. To determine how well the STRF models could account for SSA, SI was calculated for model predictions, also using Eq. 13. We then assessed the accuracy of SI predicted by STRFs in two ways: First we computed the correlation coefficient between actual and predicted SI for all the recorded cells that showed significant SI. Second, as the population mean of the squared difference between the actual and predicted SI calculated individually for each cell.

### Behavior-dependent STRFs

To measure effects of behavioral state on spectro-temporal coding, we estimated *behavior-dependent STRFs*, by allowing some or all of the fit parameters to vary between passive and active behavioral conditions [34]. Having established the efficacy of the reweighted STP STRF for passive-listening data, analysis focused on this architecture for the behavioral data.

First, a *behavior-independent STRF* provided a baseline, for which all model parameters were fixed across behavior conditions. Second, a behavior-dependent static nonlinearity allowed parameters of the static nonlinearity (Eq. 2) to vary between behavior states but kept all other parameters fixed between conditions. Third, both the linear filter parameters and static nonlinearity (Eqs. 5, 2) were allowed to vary between behavior conditions, with reweighting and STP parameters fixed across conditions. Finally, all model parameters were allowed to vary between behavior conditions. Thus, this progression of models explored the benefit of allowing increased influence of changes in behavioral state on spectro-temporal coding.

Behavior-dependent models were fit using a sequential gradient descent algorithm in the NEMS library. All models were initially fit using a behavior-independent model. The specified behavior-dependent parameters were then allowed to vary between behavioral states in a subsequent application of gradient descent. Model performance was compared as for the passive-listening data described above. For each neuron, prediction accuracy was assessed using a validation set drawn from both active and passive conditions, which was excluded from fitting, and was always the same across all models. Significant behavioral effects were indicated by improved prediction correlation for behavior-dependent STRFs over the behavior-independent STRF. Changes in tuning were measured by comparing STRF parameters between behavior conditions.

### Nonlinear encoding models for natural sounds

Encoding of natural sounds was modeled using a similar approach as for the vocalization-modulated noise. Here we focused on two models, a baseline LN STRF and a local STP STRF. Because natural sounds contain spectral features that vary across a large number of spectral channels, a different spectral filtering process was required prior to the STP stage. This was achieved using a reduced-rank STRF, where the full spectro-temporal filter was computed from the product of a small number of spectral and temporal filters [31,51]. The input spectrogram was computed from a bank of log-spaced gammatone filters, *s*(*i,t*), with *N* = 18 spectral channels [89]. Spectral tuning was modeled with a bank of *J* weight vectors, *w*(*i,j*), each of which computed a linear weighted sum of the log-compressed input spectrogram,

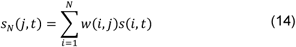

The reweighted stimuli *s*_*N*_ (*i,j*), were provided as inputs to the LN and STP STRFs (see above). Each spectral filter was initialized to have constant weights across channels. Model fitting and testing were performed using the same procedures as for the vocalization-modulated noise data (see above).

### Statistical methods

To test whether the prediction of a model for a single neuron was significantly better than chance (i.e., the model could account for any auditory response), we performed a *permutation test*. The predicted response was shuffled across time 1000 times, and the prediction correlation was calculated for each shuffle. The distribution of shuffled correlations defined a noise floor, and a *p* value was defined as the fraction of shuffled correlations greater than the correlation for the actual prediction. The Bonferroni method was used to correct for multiple comparisons when assessing significance across any of multiple models.

To compare performance of two models for a single neuron, we used a *jackknifed t-test*. The Pearson’s correlation coefficient between the actual response and response predicted by each model was calculated for 20 jackknife resamples. We then calculated the mean and standard error on the mean from the jackknifed measures [90]. The prediction of two models was considered significantly different at *p*<0.05 if the difference of the means was greater than the sum of the standard errors.

To test whether the calculated SSA Index (SI) was significantly different than chance, we performed a permutation test in which the identity of tones (standards, oddball) was shuffled, and the SI was calculated 1000 times. The real SI value was then compared to the noise floor distributions. Finally, for comparing model performance across collections of neurons, we performed a Wilcoxon signed rank test (*sign test*) between the median prediction correlation across neurons for each model.

## Author Contributions and Notes

M.L.E., Z.P.S. and S.V.D. designed research, M.L.E., Z.P.S. and S.V.D. performed research, S.V.D. wrote software, M.L.E., Z.P.S. and S.V.D. analyzed data; and M.L.E. and S.V.D. wrote the paper.

The authors declare no conflict of interest.

## Acknowledgments

This work was supported by grants from the National Institutes of Health (R01 DC014950, F31 DC016204), the Defense Advanced Research Projects Agency (D15AP00101), and a fellowship from the ARCS Foundation Oregon Chapter. The authors would like to thank Henry Cooney, Sean Slee, and Daniela Saderi for assistance with behavioral training and neurophysiological recording.

